# An automated high-throughput system for phenotypic screening of chemical libraries on *C. elegans* and parasitic nematodes

**DOI:** 10.1101/187427

**Authors:** Frederick A. Partridge, Anwen E. Brown, Steven D. Buckingham, Nicky J. Willis, Graham M. Wynne, Ruth Forman, Kathryn J. Else, Alison A. Morrison, Jacqueline B. Matthews, Angela J. Russell, David A. Lomas, David B. Sattelle

## Abstract

Parasitic nematodes infect hundreds of millions of people and farmed livestock. Further, plant parasitic nematodes result in major crop damage. The pipeline of therapeutic compounds is limited and parasite resistance to the existing anthelmintic compounds is a global threat. We have developed an INVertebrate Automated Phenotyping Platform (INVAPP) for high-throughput, plate-based chemical screening, and an algorithm (Paragon) which allows screening for compounds that have an effect on motility and development of parasitic worms. We have validated its utility by determining the efficacy of a panel of known anthelmintics against model and parasitic nematodes: *Caenorhabditis elegans, Haemonchus contortus, Teladorsagia circumcincta*, and *Trichuris muris*. We then applied the system to screen the Pathogen Box chemical library in a blinded fashion and identified known anthelmintics, including tolfenpyrad, auranofin, and mebendazole and 14 compounds previously undescribed as anthelmintics, including benzoxaborole and isoxazole chemotypes. This system offers an effective, high-throughput system for the discovery of novel anthelmintics.

## 1. Introduction

Parasitic nematodes infect around one billion people, with the soil transmitted nematodes Ascaris, hookworm and whipworm (*Trichuris trichiura*) each afflicting hundreds of millions of people (Hotez, 2013). These diseases cause high morbidity and are closely linked to poverty in the developing world. The global impact of parasitic nematodes is worsened by their effect on livestock, equids and companion animals. Parasitic nematodes of livestock are thought to cost approximately $10 billion annually with anthelmintic products comprising a major segment of the veterinary pharmaceuticals market (Roeber et al., 2013).

There are concerns that mass drug administration (MDA) programs aimed at controlling or eradicating human parasitic nematodes will become unsustainable by anthelmintic resistance, jeopardising attempts to eradicate these important tropical diseases (Prichard et al., 2012). In the case of *Trichuris*, modelling indicates that existing benzimidazole drugs are insufficiently effective for single dose MDA programs designed to break transmission; therefore eradication cannot be achieved (Keiser J and Utzinger J, 2008; Levecke et al., 2014; Turner et al., 2016). The current arsenal of veterinary anthelmintics is threatened by the emergence and spread of resistance in parasitic nematodes of domesticated animals, a recent example of which is the report of resistance to monepantel in sheep nematode species only four years after introduction of the compound (Scott et al., 2013). Multi-class resistance is also common in parasitic nematodes of ruminants (Rose et al., 2015; Sutherland, 2015) and equids (Matthews, 2014). Thus, there is an urgent requirement for the identification of novel classes of anthelmintics.

Phenotypic screening is enjoying a resurgence in recent years, and is playing an increasingly important role in the modern drug discovery paradigm (Al-Ali, 2016). Furthermore, employment of this approach using live nematodes *ex vivo* has been historically successful for target identification and lead optimisation. For example, a series of acetonitrile derivatives were identified using a *Haemonchus. contortus in vitro* larval development assay, and this led to the discovery of monepantel (Kaminsky et al., 2008).

### 1.1. Automated systems for phenotypic screening of parasitic nematodes and models of parasites

Given the need for new anthelmintics, there has been growing interest both in phenotypic screens for identification of new classes of active small molecules and in systems for quantifying relevant readouts of nematode viability such as growth and motility (Buckingham et al., 2014; Buckingham and Sattelle, 2008). Manual scoring of motility, growth or viability has been effective and used to screen libraries of up to 67,000 compounds (Burns et al., 2015; Tritten et al., 2011, 2012). Automated systems offer the potential of higher throughput and greater reliability. Such approaches include indirect assessment of viability by using the xCELLigence System; assessment of metabolic activity via colorimetric assays such as resazurin, MTT, and acid phosphatase activity; assessment of motor activity via isothermal microcalorimetry and quantification of movement-related light scattering (Nutting et al., 2015; Silbereisen et al., 2011; Smout et al., 2010; Wangchuk et al., 2016a, 2016b).

Imaging-based systems for quantification of motility or growth have also been developed. The “WormAssay” system quantifies the motility of macroscopic parasites such as *Brugia malayi* adult worms (Marcellino et al., 2012). This system has been further developed into “The Worminator”, which quantifies smaller, microscopic nematode stages and has been validated by quantifying the activity of several anthelmintics (Storey et al., 2014). This system has a reported scan time of 30 seconds per well, hence a throughput of around one and a quarter 6-well plates per hour. A system based on single-well imaging and thresholding of motile pixels with a throughput of around five 96-well plates per hour has been reported (Preston et al., 2016a). Its utility has been demonstrated by the successful screening of a 522-compound kinase inhibitor library and the 400-compound Medicine for Malaria Venture Pathogen box on *Haemonchus contortus* larvae (Preston et al., 2015, 2016b). A notable recently-described screen of the effects of 26000 compounds on *Caenorhabditis elegans* growth/survival used WormScan, a system that uses a conventional flat-bed scanner to capture two frames of images of whole plates and then uses an algorithm based on the difference image to assign a value to each well that reflects motility/growth (Mathew et al., 2012, 2016). This led to the identification of several compounds with previously unreported anthelmintic activity, including compounds targeting PINK-1 and MEV-1. The authors reported a throughput of approximately 25-40 96-well plates per hour.

### 1.2. Developing a new robust motility/growth quantification system focussed on rapid, high-throughput chemical screening

It is clear that recent developments in phenotypic screening of parasitic and model nematodes have led to an acceleration of the discovery of potential novel anthelmintic compounds. Given the large sizes of drug-like compound libraries and the need to efficiently identify the hit compounds therein that are have the potential to be developed into potent and selective anthelmintic lead molecules, it is desirable that nematode phenotypic screening be further accelerated. Here, we present the development of the INVAPP / Paragon system, which quantifies nematode motility/growth with a throughput of approximately 100 96-well plates per hour, with a robust and unbiased approach. We validate the system by quantifying the activity of a panel of known anthelmintics on a variety of nematode species, and then by screening, in a blinded fashion, the Medicines for Malaria Venture Pathogen Box for compounds that block or reduce nematode growth.

## 2. Materials and methods

### 2.1. Ethics statement

All mouse experiments were approved by the University of Manchester Animal Welfare and Ethical Review Board and performed under the regulation of the United Kingdom Home Office Scientific Procedures Act (1986) and the Home Office project licence 70/8127.

All ovine experiments were approved by the Moredun Research Institute Experiments and Ethics Committee and performed under the regulation of the United Kingdom Home Office Scientific Procedures Act (1986) and the Home Office project licence 60/04421.

### 2.2. INVAPP / Paragon system

The INVAPP / Paragon system consists of a fast high-resolution camera (Andor Neo, resolution 2560x2160, maximum frame rate 100 frames per second) with a line-scan lens (Pentax YF3528). Microtiter plates are placed in a holder built into the cabinet and imaged from below. Illumination is provided by an LED panel with acrylic diffuser. Movies were captured using μManager (Edelstein et al., 2014). The desirable movie frame length and duration of filming depends on the particular organism under study and is specified below. Movies were analysed using MATLAB scripts. Briefly, movies were analysed by calculating the variance through time for each pixel. The distribution of these pixel variances was then considered, and pixels whose variance was above the threshold (typically those greater than one standard deviation away from the mean variance) were considered ‘motile’. Motile pixels were then counted and assigned by well, generating a movement score for each well. The source code for this software has been released under the open source MIT license and is available at https://github.com/fpartridge/invapp-paragon. A further MATLAB script has been provided for batch processing of movies.

### 2.3. Caenorhabditis elegans motility and growth assays

*C. elegans* strains were maintained at 20 °C on nematode growth medium agar seeded with the *E. coli* strain OP50. To obtain worms for screening, a mixed-stage liquid culture was prepared by washing well-fed worms from one small NGM plate into a medium of 50 ml S-complete buffer with a pellet of approximately 2-3 g *E. coli* HB101. Cultures were agitated at 200 rpm, 20 °C, until there were many adults present, then synchronised at the L1 stage by bleaching. Fifty millilitre cultures were pelleted and bleaching mix (1.5 ml 4M NaOH, 2.4 ml NaOCl, 2.1 ml water) added. Mixing for 4 minutes led to the release of embryos, which were washed three times with 50 ml S-basal medium. The cultures were incubated in 50 ml S-basal at 20 °C and agitated at 200 rpm overnight to allow eggs to hatch and arrest as a synchronous L1 population.

For the growth assay, *C. elegans* were cultured in a 96-well plate format. Synchronised L1 were diluted to approximately 20 worms per 50 μl in S complete medium with around 1% w/v HB101 *E. coli*. Assay plates were prepared with 49 μl of S-basal and 1μl of DMSO or compound in DMSO solution per well. Next, 50 μl of the L1 suspension were added to each well. Plates were incubated at 20°C before imaging using the INVAPP / Paragon system 5 days later. Prior to imaging, worm motion was stimulated mechanically by inserting and removing a 96-well PCR plate into/from the wells of the assay plate. Whole-plate 200 frame movies were recorded at 30 frames per second (7 seconds total).

For the adult motility assay, synchronised L1 were refed as a bulk 50 ml culture and cultured at 20 °C until they developed into young adults. Worms were washed in S-basal and dispensed, approximately 20 worms per well, into 96-well plates with compound dissolved in DMSO, or DMSO alone and then incubated for 3 hours. Whole-plate 200 frame movies were recorded at 30 frames per second (7 seconds total).

### 2.4. Trichuris muris motility assay For the adult

*T. muris* motility assay, male and female severe combined immune deficiency mice (bred in the Biological Services Facility at the University of Manchester) were infected with 200 embryonated *T. muris* eggs in water by oral gavage. After 35 days, mice were killed and their caecae and colons removed, opened longitudinally, and washed with pre-warmed Roswell Park Memorial Institute (RPMI) 1640 media supplemented with penicillin (500 U/ml) and streptomycin (500 μg/ml). Adult *T. muris* worms were then removed using fine forceps and maintained in RPMI-1640/penicillin/streptomycin media at approximately 37 °C and studied on the same day. Individual live worms were placed into 96 well plates containing 75 μl of RPMI-1640/penicillin/streptomycin medium plus 1% v/v final concentration of DMSO or compound dissolved in DMSO. Plates were incubated at 37 °C, 5% v/v CO_2_, and motility was analysed after 24 hours. Whole plate 200 frame movies were recorded at 10 frames per second (20 seconds total).

### 2.5. Ovine parasitic nematode isolation

Six month-old, male Texel cross lambs that were raised under helminth-free conditions were infected *per os* either with 15,000 *T. circumcincta* (isolate MTci2) larvae (L_3_) or 5,000 *H. contortus* (isolate MHco3) L_3_. Once a patent infection was detected by the observation of nematodes eggs in faeces (around 21 days post-infection), the lambs were fitted with a collection harness and bag to enable faecal collection. Pelleted faeces were collected from each bag 24 hours later and placed in a covered seed tray, which was incubated at room temperature (>15 °C) for 10 days before larval recovery using the modified Baermann technique (*Manual of veterinary parasitological laboratory techniques.*, 1986). The next day L_3_ were recovered in ~250 ml H_2_O, number estimation performed then the larvae were stored in 100 ml tap water in 75 cm^2^ surface area vented cap, suspension culture flasks (Sarstedt Ltd UK) at ~ 5 °C for MTci2 and 8 °C for MHco3, respectively.

### 2.6. T. circumcincta assay

Ensheathed L3 *T. circumcincta* were used in this assay. Approximately 30 worms were added to wells containing compound + DMSO or DMSO alone (final concentration 1% v/v). Worms were incubated for 2 hours in the dark at 25 °C. Movement was stimulated by illuminating the plate with bright white light for 3 minutes (Zeiss HAL100), before acquiring movies on the INVAPP / Paragon system (200 frames, 10 frames per second).

### 2.7. H. contortus assay

*H. contortus* exsheathed L3 (xL3) were prepared by treatment with sodium hypochlorite (Preston et al., 2015). A 1ml solution containing around 1000 *H. contortus* L3 was placed in a 35 mm petri dish. 20 μl sodium hypochlorite solution (Fisher, S/5040/PC17) was added and incubated at room temperature for 4 minutes. Exsheathment was monitored using a dissecting microscope. The worms were filtered using a 10 μm cell strainer (pluriSelect), rinsed with 10x 1 ml S-basal solution, and eluted with 1 ml S-basal solution. Around 30 xL3 worms (in S-basal solution) were added to wells containing compound + DMSO or DMSO alone (final DMSO concentration 1% v/v). Worms were incubated for 2 hours in the dark at 25 °C. Movement was stimulated by illuminating the plate with bright white light for 3 minutes (Zeiss HAL100), before acquiring movies on the INVAPP / Paragon system (200 frames, 30 frames per second). The INVAPP / Paragon *movementIndexThreshold* parameter was set to 2 for analysis of *H. contortus xL3*, due to a lower prior expectation of worm movement in the movie for this nematode.

### 2.8. Pathogen Box screening

The Pathogen Box library was obtained from the Medicines for Malaria Venture as 10 mM solutions in DMSO, and then diluted in DMSO to 1 mM. It was then screened in the *C. elegans* growth assay as described (final concentration 10 μM, n=5, 1% v/v final DMSO). Solid material for confirmatory screening of actives was obtained from Sigma-Aldrich (tolfenpyrad) and Santa Cruz Biotechnology (auranofin). Solid samples of MMV007920, MMV020152, MMV652003 and MMV688372 were obtained from the Medicines for Malaria Venture.

## 3. Results

### 3.1. INVAPP / Paragon: a high-throughput assay for quantifying nematode motility and growth

We wanted to develop an assay for screening small molecules for their effect on the motility and growth of diverse parasites. A high-throughput and automated system was particularly desirable, given the large size of available small molecule libraries. To this end, we designed the INVAPP / Paragon system. A schematic of the INVAPP hardware is shown in Fig 1A. This allows recording of movies of entire microplates at high frame rate, reducing the per plate acquisition time to 10-30 seconds. Tens of thousands of compounds or conditions can therefore be readily screened per day.

**Fig 1.**
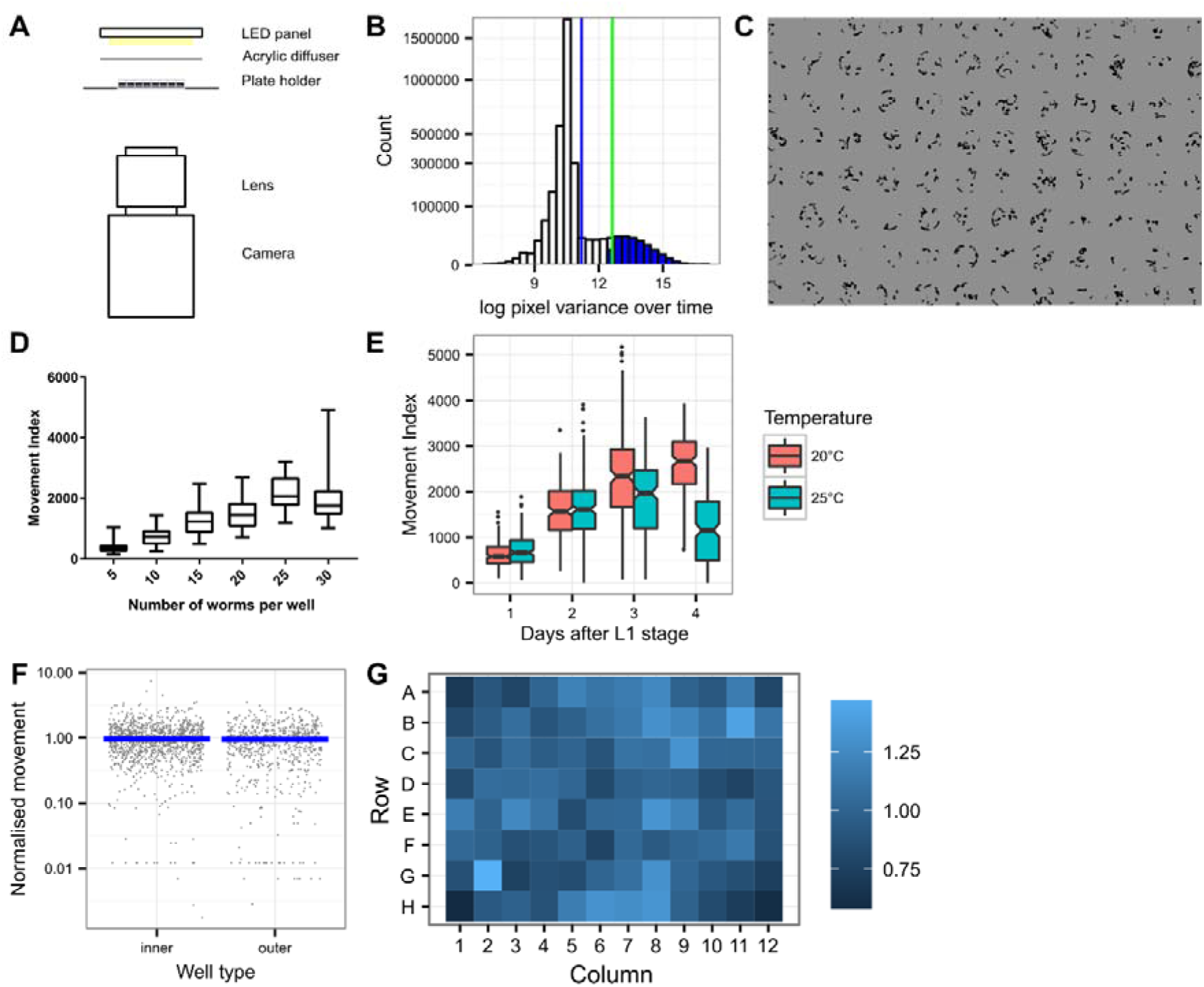
The INVAPP / Paragon movement index algorithm is a fully-automated high-throughput system able to determine motility and growth rate. (A) Schematic of the INVAPP setup (B) Principle of the algorithm: thresholding of moving pixels by statistical analysis of variance of each pixel through time. Histogram shows the distribution of pixel variance over time. Blue vertical line indicates mean pixel variance. The green vertical line indicates mean plus standard deviation of pixel variance; the blue shaded portion of the histogram indicates pixels that exceed this threshold so are deemed to be motile. (C) Image of 96-well plate containing *C. elegans* adults processed by the INVAPP / Paragon movement index system. Dark pixels are those categorized as moving by the algorithm. (D) Increasing the number of *C. elegans* worms per well leads to increase in reported movement index. Boxplot bars indicate 95% confidence interval. (E) Movement index algorithm is able to quantify *C. elegans* growth in 96-well plates. Movement index increases with growth. Synchronized L1 population refed on day 0. Decrease in movement index in 25°C group on Day 4 reflects completion of the *C. elegans* lifecycle and exhaustion of the bacterial food source. Boxplot notches indicate 95% confidence interval, n=192. (F) Absence of edge effects in this assay – analysis of a 1920-well *C. elegans* growth dataset shows no difference of the normalised movement score for 96-well plate outer edge wells (the wells found in columns 1 and 12 or rows A and H) compared to the score for inner wells (the other wells in the plate). Movement index for each well is normalised by dividing by the mean movement index for all wells of that plate. The blue bar indicates median. (G) No edge effects or other inhomogeneity across the plate – heat map shows average normalised movement index for each well location.

We took a statistical approach to quantifying motility. The variance through time for each pixel in the plate was calculated and the distribution of the variances examined. Pixels whose variance is greater than a threshold of the mean plus typically one standard deviation are determined to be “motile” (Fig 1B). For organisms that are particularly small or have limited motility, such as *H. contortus* L3s, this threshold was increased to the mean plus two standard deviations, reflecting a lower prior expectation of worm movement in the movie. A similar approach was recently reported for the determination of *H. contortus* motility (Preston et al., 2015). An example of this thresholding model is shown in Fig 1C, which shows analysis of a 96-well plate containing adult *C. elegans*. Dark pixels are those that have been determined to be motile. Once the motility threshold has been applied to the data, ‘motile’ pixels are assigned by well to their plate location and counted. All analysis is fully automated via a set of MATLAB scripts, available at https://github.com/fpartridge/invapp-paragon.

This approach was able to determine motility. To illustrate this, we analysed plates containing a variable number of synchronized adult *C. elegans* worms. As expected, quantified movement increased with the number of worms per well, reflecting a larger number of ‘motile’ pixels in the recording (Fig 1D).

The system was also able to quantify nematode growth. To test this, we synchronised *C. elegans* at the L1 stage, before refeeding them in plates at two temperatures commonly used in *C. elegans* culture (20 °C and 25 °C). Plates were then analysed using INVAPP / Paragon every 24 hours. The results are shown in Fig 1E. The quantified movement index increases as worms develop from L1 to adult stage. The drop in motility in the 25 °C group on day 5 reflects growth of L1 progeny leading to exhaustion of the bacterial food source and starvation.

When establishing a high-throughput assay it is important to consider the issue of edge effects (Malo et al., 2006). Systematic biases across the plate are particularly common around edges. Typical causes are evaporation, which is often worse at the edges, or temperature inhomogeneity. In our assay, given that it involves imaging of whole plates, we also wanted to exclude the possibility of systemic bias caused by optical distortion. To address these concerns, we analysed a 1920-well *C. elegans* growth dataset. This was chosen because the long four-day incubation time gave the maximum possibility of confounding evaporation differences. We classified wells on the outer rows and columns of the plate as being outer wells, and compared their quantified motility to the inner wells (Fig 1F). There was no significant difference between these groups (Mann-Whitney-Wilcoxon test, P=0.77), and therefore no evidence of problematic edge effects in this assay. To further exclude the possibility of assay inhomogeneity across the plate, we calculated a heat map showing average normalised motility for each well (Fig 1G). Again, this showed no evidence of systemic bias by plate position.

### 3.2. Validation of the INVAPP / Paragon assay using existing commercial anthelmintic standards

Having set up this high-throughput motility and growth assay, we wanted to validate its utility by examining the effects of a panel of known anthelmintics. We selected nine anthelmintics with a variety of reported mechanisms of action. Piperazine is a GABA agonist that acts at the neuromuscular junction (Martin, 1985). Diethylcarbamazine has been proposed to have a similar mechanism, although other mechanisms including targeting host arachidonic acid metabolism are also thought to be important (Maizels and Denham, 1992). Levamisole, oxantel and pyrantel are nicotinic acetylcholine receptor agonists that induce spastic paralysis (Martin et al., 1997). Mebendazole is an inhibitor of beta-tubulin polymerisation (Driscoll et al., 1989). Ivermectin is a positive allosteric modulator of glutamate-gated chloride channels although other targets have also been suggested (Cully et al., 1994). Trichlorfon is a member of the organophosphate group of acetylcholine esterase inhibitors. Praziquantel is thought to act by disrupting calcium ion homeostasis but its target is unclear (Caffrey, 2007)

We first measured concentration-response curves for this panel of anthelmintics in an acute treatment (3 hour) adult *C. elegans* assay. The results are shown in Fig 2. As expected, the major ion channel-targeting drugs ivermectin, levamisole, oxantel and pyrantel were active in this assay, reflecting their direct effects on worm motility.

**Fig 2.**
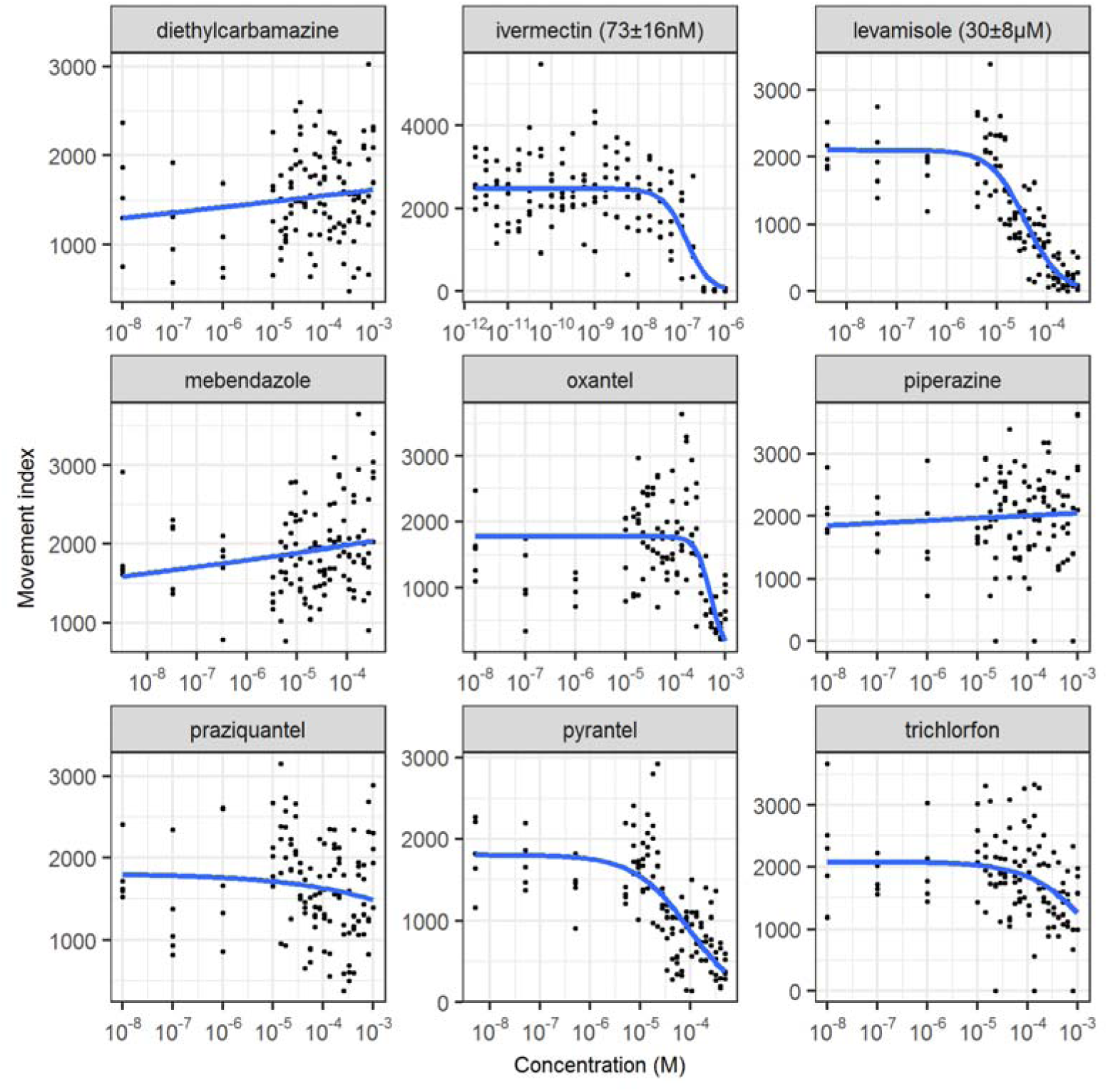
Determination of the effect of acute treatment with important anthelmintics on *C. elegans* adult motility using the INVAPP / Paragon movement index algorithm. Blue line fitted using the 3-parameter log-logistic model. EC_50_ values are shown in parentheses, with standard error for this estimate calculated using *drc* (Ritz and Streibig, 2005).

We then measured concentration-response curves for this panel of anthelmintics in a chronic treatment *C. elegans* growth assay. The results are shown in Fig 3. Activity was again found with ivermectin, levamisole, oxantel and pyrantel and, additionally, with mebendazole, piperazine and trichlorfon. This reflects that some anthelmintic modes of action may not be measured in purely acute motility assays, supporting the use of assays that involve growth or development. Diethylcarbamazine and praziquantel were not active in these assay as expected. This reflects previously reported low *in vitro* activity of diethylcarbamazine and its proposed mechanism of acting on host arachidonic acid metabolism (Maizels and Denham, 1992). Praziquantel, used primarily to treat flukes and tapeworms, is known to have limited efficacy against nematodes (Holden-Dye, 2007). Successfully demonstrating the ability of the INVAPP / Paragon system to determine the effects of known anthelmintics increased our confidence in this approach.

**Fig 3.**
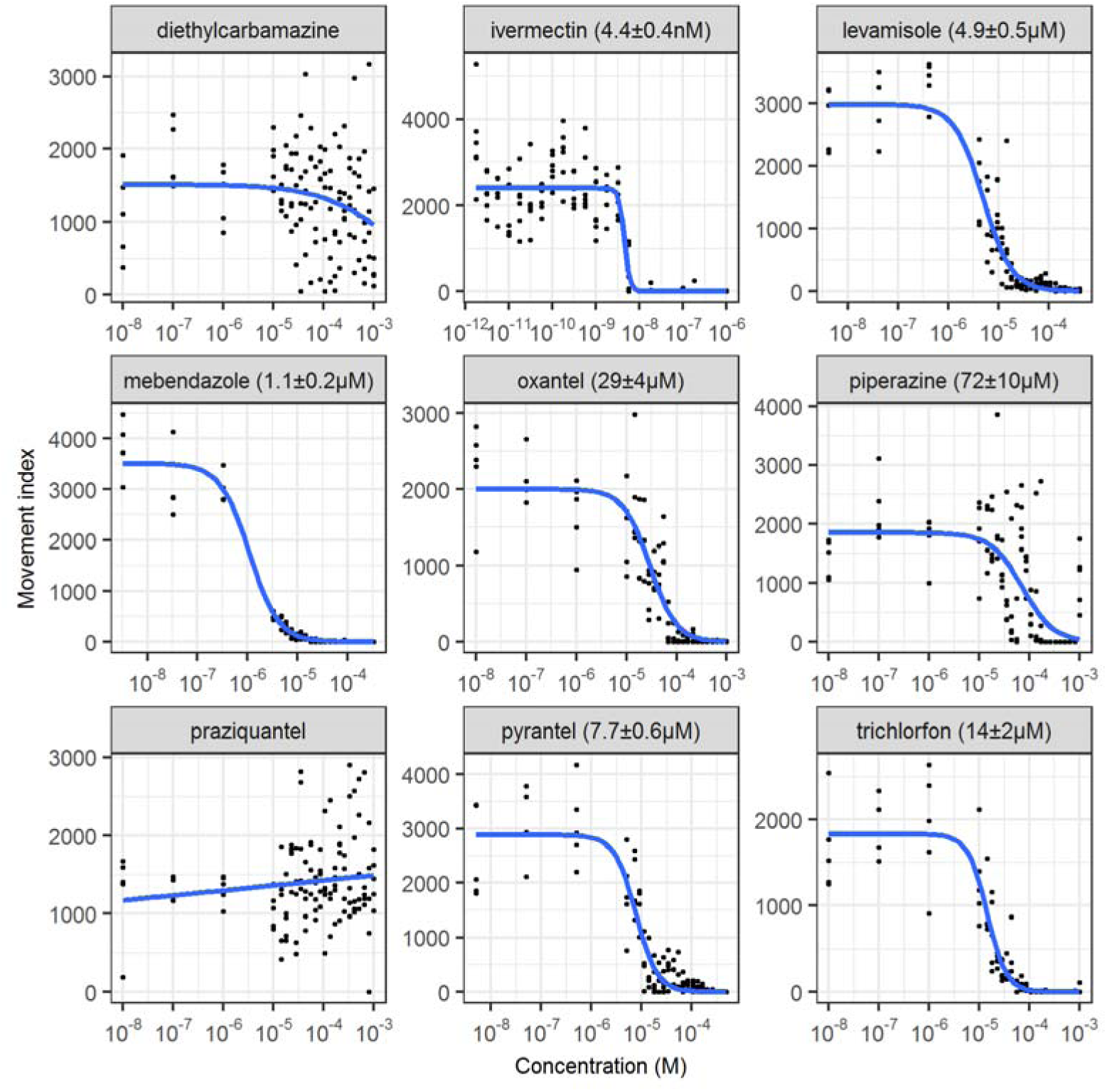
Determination of the effect of important anthelmintics on *C. elegans* growth using the INVAPP / Paragon movement index algorithm. Blue line fitted using the 3-parameter log-logistic model. EC_50_ values are shown in parentheses, with standard error for this estimate calculated using *drc* (Ritz and Streibig, 2005).

### 3.3. Adaptation of the INVAPP / Paragon assay to parasitic nematodes

*C. elegans* is a useful model because of the ease of laboratory culture and access to powerful genetic tools. However, it is also valuable to screen parasitic nematodes directly. *T. circumcincta* is a globally important parasitic nematode that infects small ruminants. We tested the ability of the INVAPP / Paragon system to quantify the activity of the anthelmintic ivermectin on ensheathed *T. circumcincta* L3. Worms were incubated at 25 °C with the compounds for 2 hours in the dark, after which movement was induced by bright light and movies recorded on the INVAPP system. Fig 4A shows an example of the INVAPP / Paragon motility thresholding of single wells treated with DMSO or DMSO plus 100 μM ivermectin, showing much reduced, but not abolished movement, of these larvae in the assay. We obtained a concentration-response curve for ivermectin (Fig 4B), demonstrating the ability of the system to quantify motility of this parasite.

**Fig 4.**
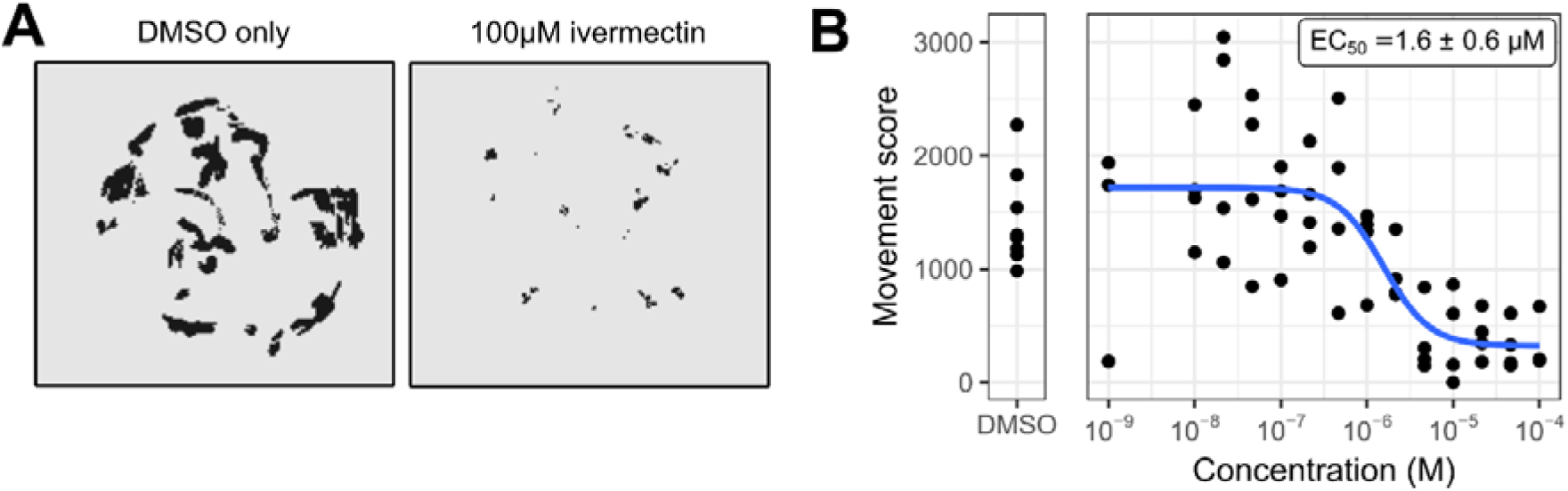
Utilising the INVAPP / Paragon system with ensheathed L3 *T. circumcincta*. (A) INVAPP / Paragon motility thresholding of single wells treated with DMSO or a DMSO solution of ivermectin (assay concentration 100 μM). Black pixels are scored as motile. (B) Concentration-response curve for treatment with the anthelmintic ivermectin, n=4. Curve fitted using the four-parameter log-logistic function. Error range on the EC_50_ estimate indicates standard error (delta method).

*H. contortus*, the barber’s pole worm, is another economically important gastrointestinal parasite of small ruminants. We tested the ability of the INVAPP / Paragon system to quantify the activity of ivermectin on exsheathed L3 (xL3) *H. contortus.* Worms were incubated at 25 °C with the compounds for 2 hours in the dark, after which movement was induced by bright light and movies recorded on the INVAPP system. An example of the INVAPP / Paragon motility thresholding of single wells treated with DMSO or a DMSO solution of ivermectin (assay concentration 100 μM) is shown in Fig 5A. Movement of ivermectin-treated xL3s in the assay was considerably reduced. A concentration-response curve for ivermectin is shown in Fig 5B, illustrating the ability of the system to quantify motility of this parasite.

**Fig 5.**
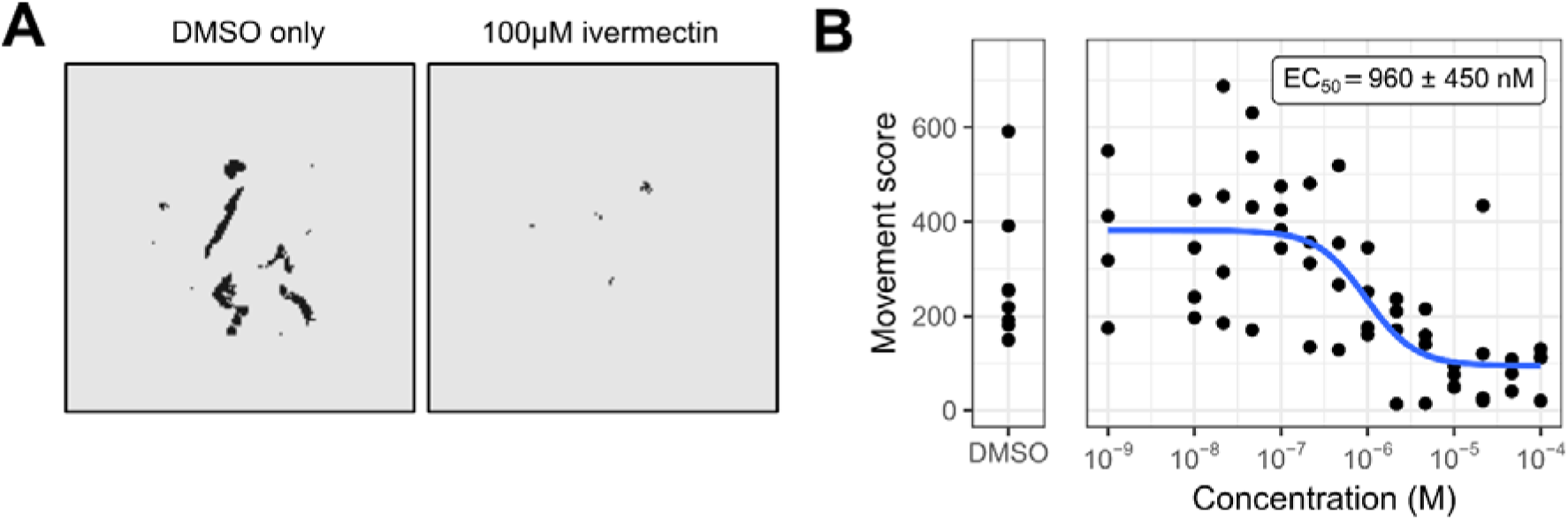
Utilising the INVAPP / Paragon system with exsheathed L3 *H. contortus*. (A) INVAPP / Paragon motility thresholding of single wells treated with DMSO or a DMSO solution of ivermectin (assay concentration 100 μM). Black pixels are scored as motile. (B) Concentration-response curve for treatment with the anthelmintic ivermectin, n=4. Curve fitted using the four-parameter log-logistic function. Error range on the EC_50_ estimate indicates standard error (delta method).

*Trichuris muris*, which infects mice, is a widely used laboratory model for investigating trichuriasis and has been used as a system for *ex vivo* and *in vivo* testing of anthelmintic candidates for activity against whipworm (Hurst et al., 2014; Tritten et al., 2011; Wimmersberger et al., 2013). We tested the ability of the INVAPP / Paragon system to quantify the activity of the anthelmintic levamisole on adult *T. muris*. Worms were incubated at 37 °C with the compounds for 24 hours in the dark, after which movement was recorded on the INVAPP system. Fig 6A shows control and levamisole treated wells. For illustration, selected frames of the movie are shown (3-second intervals). The readout of ‘motile’ pixels as determined by the INVAPP / Paragon algorithm is shown in Fig 6B, showing the quantification of the loss of motility of the levamisole-treated worm. A concentration-response curve for levamisole was measured (Fig 6C), which illustrates the ability of the system to quantify motility of this parasite and thus measure anthelmintic activity.

**Fig 6.**
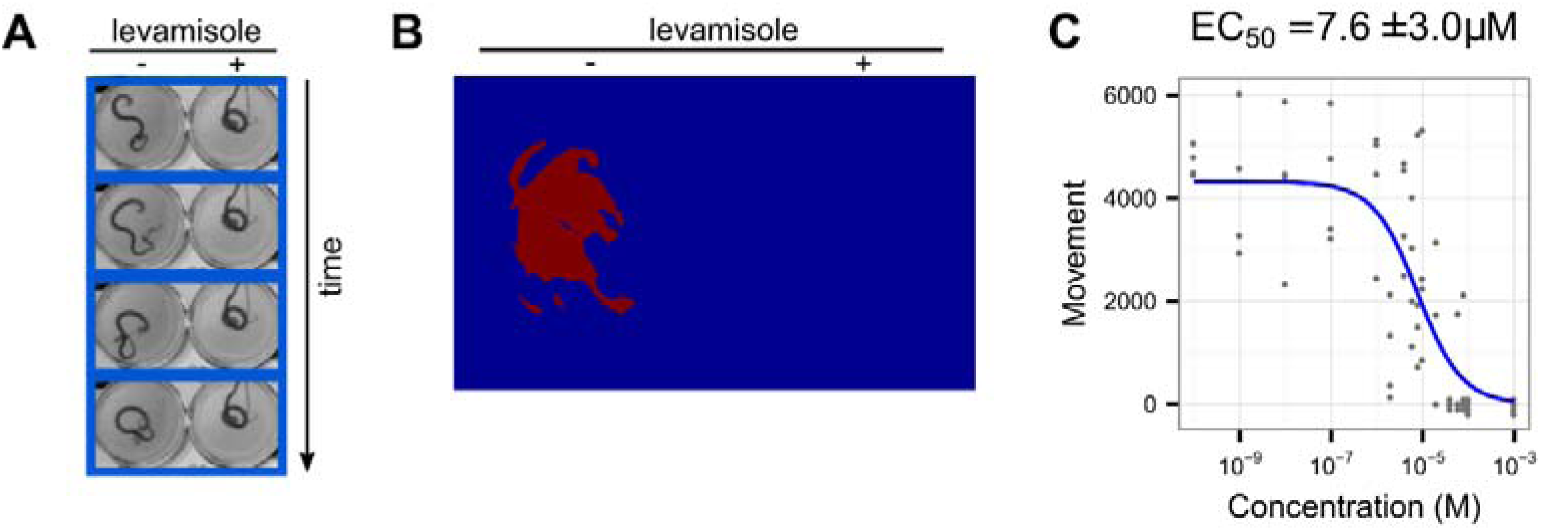
Utilising the INVAPP / Paragon system with adult *T. muris*. (A) Example of section of movie, showing control and levamisole-treated wells. Selected movie frames are shown at three-second intervals (B) Pixels for these wells that are considered to be motile as determined by the algorithm are shown in red. (C) Concentration-response curves for treatment with the anthelmintic levamisole, n=4. Curve fitted using the three-parameter log-logistic function. Error range on the EC_50_ estimate indicates standard error (delta method).

### 3.4. Screening the Pathogen Box for compounds that affect C. elegans growth

We then applied the INVAPP / Paragon assay system to the identification of novel anthelmintic small molecules. We used the Pathogen Box, a collection of 400 diverse drug-like molecules that are known to be active against various neglected disease pathogens (particularly, tuberculosis (or *Mycobacterium* spp.), malaria and kinetoplastid protozoa). This library is distributed as an open-science project by the Medicines for Malaria Venture. A screen of this library for compounds affecting motility of exsheathed L3 of *H. contortus* has been published recently (Preston et al., 2016b), which identified the insecticide, tolfenpyrad, as active against the larvae. To complement this approach, and with the aim of identifying compounds blocking growth of nematodes as opposed to solely immobilising them, we screened the library in a blinded fashion (n=5, concentration 10 μM) using the INVAPP / Paragon *C. elegans* growth assay. For each compound, motility change and significance (Mann-Whitney-Wilcoxon test) were calculated relative to DMSO-only control wells. A volcano plot showing the results of this screen is shown in Fig 7A. Compounds that reproducibly reduced growth are found towards the top left of this plot. To confirm identity of the hit molecules, we retested the top 20 putative hit compounds with the same library material (n=5, concentration 10 μM) in the same INVAPP / Paragon *C. elegans* growth assay (Fig 7B). A total of 18 out of 20 compounds were active (P < 0.05, Dunnett’s multiple comparison test).

The combined assay results from the primary and secondary screen are found in Supplementary File 1 and have been recorded in the PubChem database with Assay ID 1259336.

**Fig 7.**
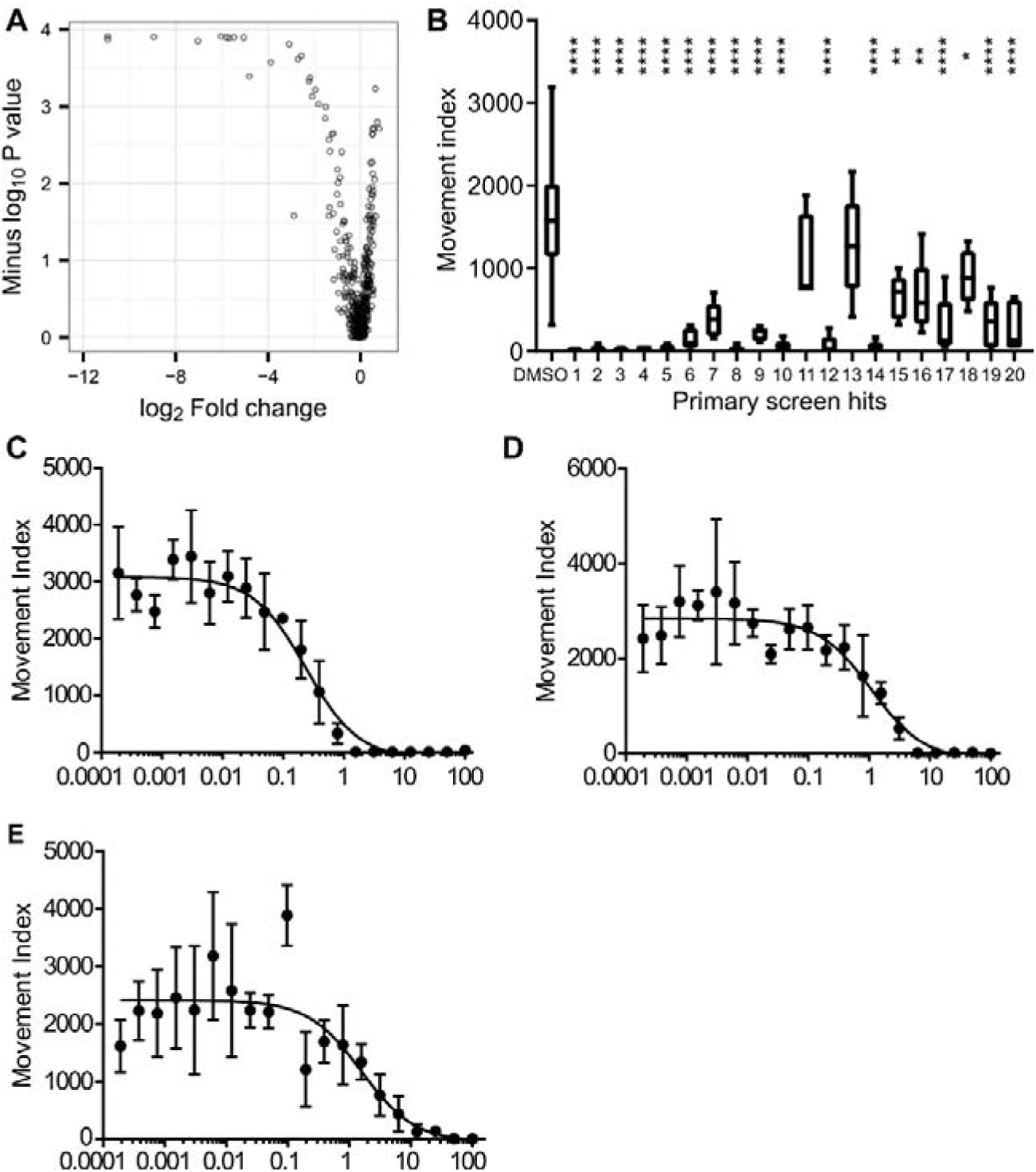
Screening the Pathogen Box in the *C. elegans* growth screen. (A) Volcano plot showing the results of the primary screen (n=5, concentration = 10 μM). Each point represents one compound. Effective size is shown on the x axis, as log_2_-fold change (ratio of the median movement for the repeats of the compound to the median movement of DMSO-only wells). Statistical significance is shown on the y axis as the-log_10_ P value in the Mann-Whitney-Wilcoxon test. A location at the top left of this plot indicates anthelmintic activity. (B) Secondary rescreen of hit compounds, in order of their activity in the primary screen, from library material. Statistical significance compared to DMSO-only control calculated by Dunnett’s multiple comparison test (n=5, * indicates P < 0.05, ** indicates P < 0.005, **** indicates P < 0.0005). (C,D,E) Concentration-response curves showing the activity of known anthelmintics – (C) tolfenpyrad, (D) auranofin, (E) isradipine – that were found in the Pathogen box screen retested using solid material (supplied by the Medicines for Malaria Venture) in the *C. elegans* growth assay. A concentration-response curve for mebendazole in the assay was presented in Fig. 3. Error bars indicate standard deviation, n=4. Curve fitting was undertaken using three parameter log logistic model (Graphpad Prism).

### 3.5. Identification of known anthelmintic compounds by screening the Pathogen Box in the C. elegans growth assay

We first considered known anthelmintic compounds that we found to be active in this screen (Table 1). Mebendazole, an anthelmintic from the benzimidazole group, acts by inhibiting microtubule synthesis. Tolfenpyrad is a broad-spectrum acaricide and insecticide that acts as an inhibitor of complex I of the electron transport chain. It has been recently reported to reduce motility of *H. contortus* exsheathed L3 and to block L3 to L4 development of this parasite *in vitro* (Preston et al., 2016b). We confirmed activity of this compound using solid material. As shown in Fig 7C, the EC_50_ was 200 ± 40 nM. Independently identifying these known anthelmintic compounds using a blinded screening approach further validates the INVAPP/Paragon system as a robust high throughput screening approach.

**Table 1.** Named compounds that were active in the *C. elegans* growth screen. Log_2_-fold change in growth estimate compared to DMSO-only controls. Hit ID is as in Fig 7B. EC_50_ confidence interval is the standard error. The EC_50_ estimate for mebendazole is from Fig. 3.

| Compound | MMV ID | PubChem CID | Hit ID | Log<sub>2</sub> fold change in growth |  | EC<sub>50</sub> (μM) ± standard error (solid material) |
| --- | --- | --- | --- | --- | --- | --- |
|  |  |  |  | (1° screen) | (2° screen) |  |
| Tolfenpyrad | MMV688934 | 10110536 | 1 | -10.9 | -10.6 | 0.2 ± 0.04 |
| Auranofin | MMV688978 | 24199313 | 2 | -10.9 | -7.2 | 1.1 ± 0.3 |
| Mebendazole | MMV003152 | 4030 | 5 | -6.0 | -5.7 | 1.1 ± 0.2 |
| Isradipine | MMV001493 | 158617 | 16 | -2.2 | -1.4 | 1.6 ± 0.7 |
| Tavaborole | n/a | 11499245 |  | n/a | n/a | 8.6 ± 1.9 |

Auranofin is a gold(I) compound originally developed for the treatment of rheumatoid arthritis. It has received attention for repurposing as an anti-cancer agent, with a number of clinical trials under way. It has been shown that auranofin has *in vitro* and *in vivo* activity in several models of parasitic diseases, including schistosomiasis (Kuntz et al., 2007), amoebiasis (Debnath et al., 2012), leishmaniasis (Sharlow et al., 2014) and onchocerciasis (Bulman et al., 2015). A phase IIa trial of auranofin for gastrointestinal protozoal infection is ongoing. We confirmed activity of this compound in the *C. elegans* growth assay using solid material. As shown in Fig 7D, the EC_50_ of this compound was 1.1 ± 0.3μM.

Isradipine is an antihypertensive drug that belongs to the dihydropyridine family of L-type calcium channel blockers. A structurally related dihydropyridine, nemadipine-A, has been shown to cause growth and egg laying defects in *C. elegans* by antagonising the L-type calcium channel α_1_-subunit EGL-19 (Kwok et al., 2006). We confirmed activity of this compound in the *C. elegans* growth assay using solid material. As shown in Fig 7E, the EC_50_ of this compound was 1.6 ± 0.7 μM.

### 3.6. Novel anthelmintics that block C. elegans growth

Fourteen compounds without previously-described anthelmintic activity were identified in the Pathogen Box screen (Table 2). The structures of the most active compounds are shown in Fig 8. We examined four of these compounds more closely. First, we determined activity, using solid material, in the *C. elegans* growth assay (Fig 9). The EC_50_ values for the confirmatory assay are shown in Table 2. These results have been recorded in the PubChem database with Assay ID 1259335.

**Fig 8.**
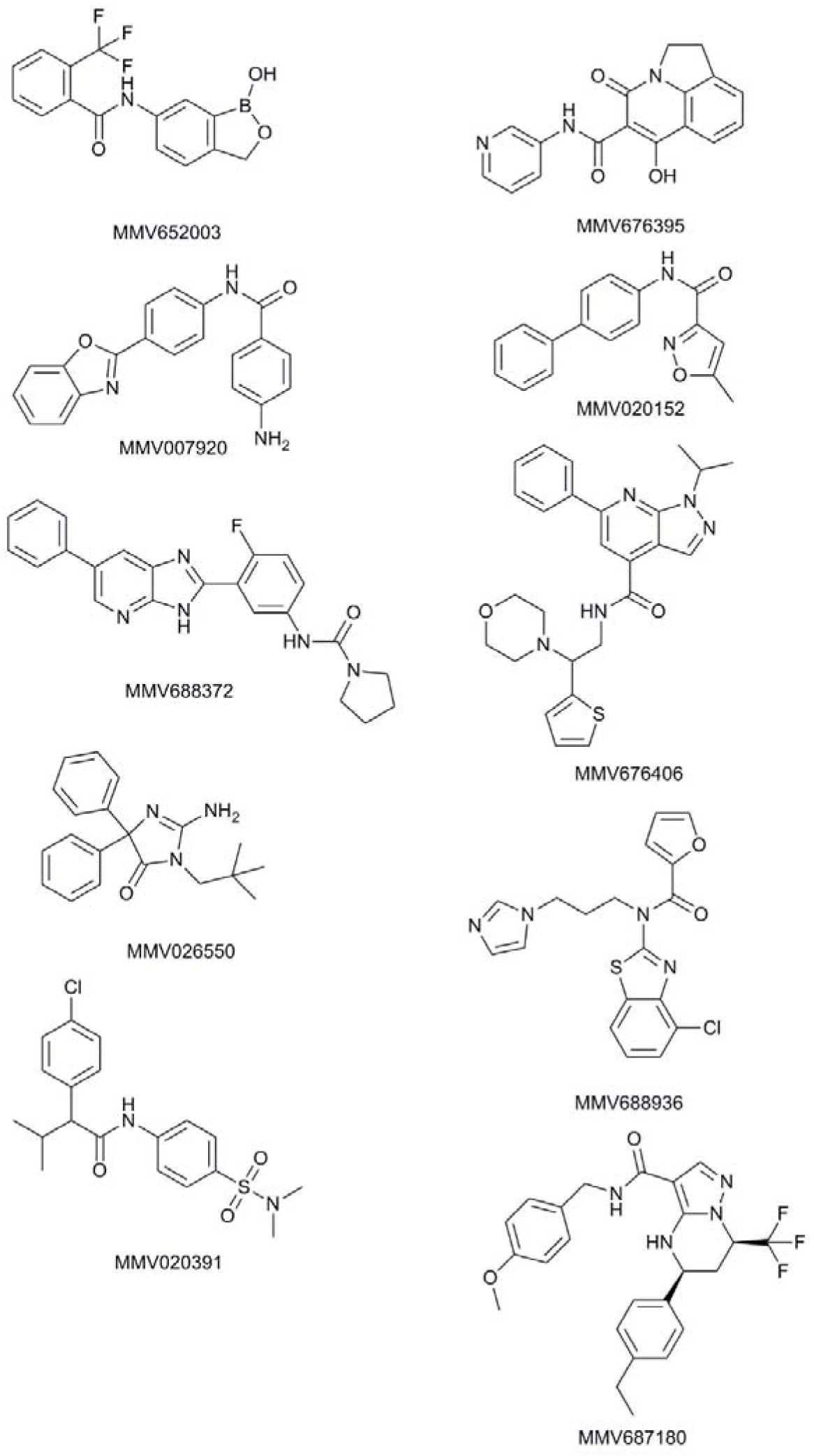
Structures of the most active compounds, previously undescribed as anthelmintics, which were found to have activity in the *C. elegans* growth screen. Compound numbers from Medicines for Malaria Venture Pathogen Box.

**Table 2.** Compounds with previously unreported anthelmintic activity that were active in the *C. elegans* growth screen. MMV ID is the compound identifier for the Medicines for Malaria Venture. PubChem CID is the compound identifier for the PubChem database. Hit ID is as in Fig 7B. Log_2_ fold change in the growth estimate in the INVAPP / Paragon assay compared to DMSO-only controls.

| MMV ID | PubChem CID | Hit ID | Log<sub>2</sub> fold change in growth |  | EC<sub>50</sub> (μM) |
| --- | --- | --- | --- | --- | --- |
|  |  |  | (1° screen) | (2° screen) |  |
| MMV652003 | 46196110 | 3 | -8.9 | -7.6 | 4.3±2.5 |
| MMV007920 | 721133 | 4 | -7.0 | -5.4 | 6.2±2.4 |
| MMV688372 | 72710598 | 6 | -5.8 | -4.0 | 3.3±1.7 |
| MMV675994 | 44222802 | 7 | -5.7 | -2.0 | n.d. |
| MMV026550 | 44530521 | 8 | -5.5 | -5.9 | n.d. |
| MMV020391 | 7918647 | 9 | -5.0 | -2.7 | n.d. |
| MMV676395 | 54678166 | 10 | -4.8 | -4.8 | n.d. |
| MMV020152 | 8880740 | 12 | -3.1 | -6.9 | 2.5±1.0 |
| MMV676406 | 30238526 | 14 | -2.7 | -8.0 | n.d. |
| MMV688417 | 58346931 | 15 | -2.6 | -1.1 | n.d. |
| MMV688936 | 18589797 | 17 | -2.2 | -3.7 | n.d. |
| MMV1028806 | 16387386 | 18 | -2.1 | -0.8 | n.d. |
| MMV688888 | 5179236 | 19 | -2.0 | -2.1 | n.d. |
| MMV687180 | 41058173 | 20 | -1.8 | -3.6 | n.d. |

**Fig 9.**
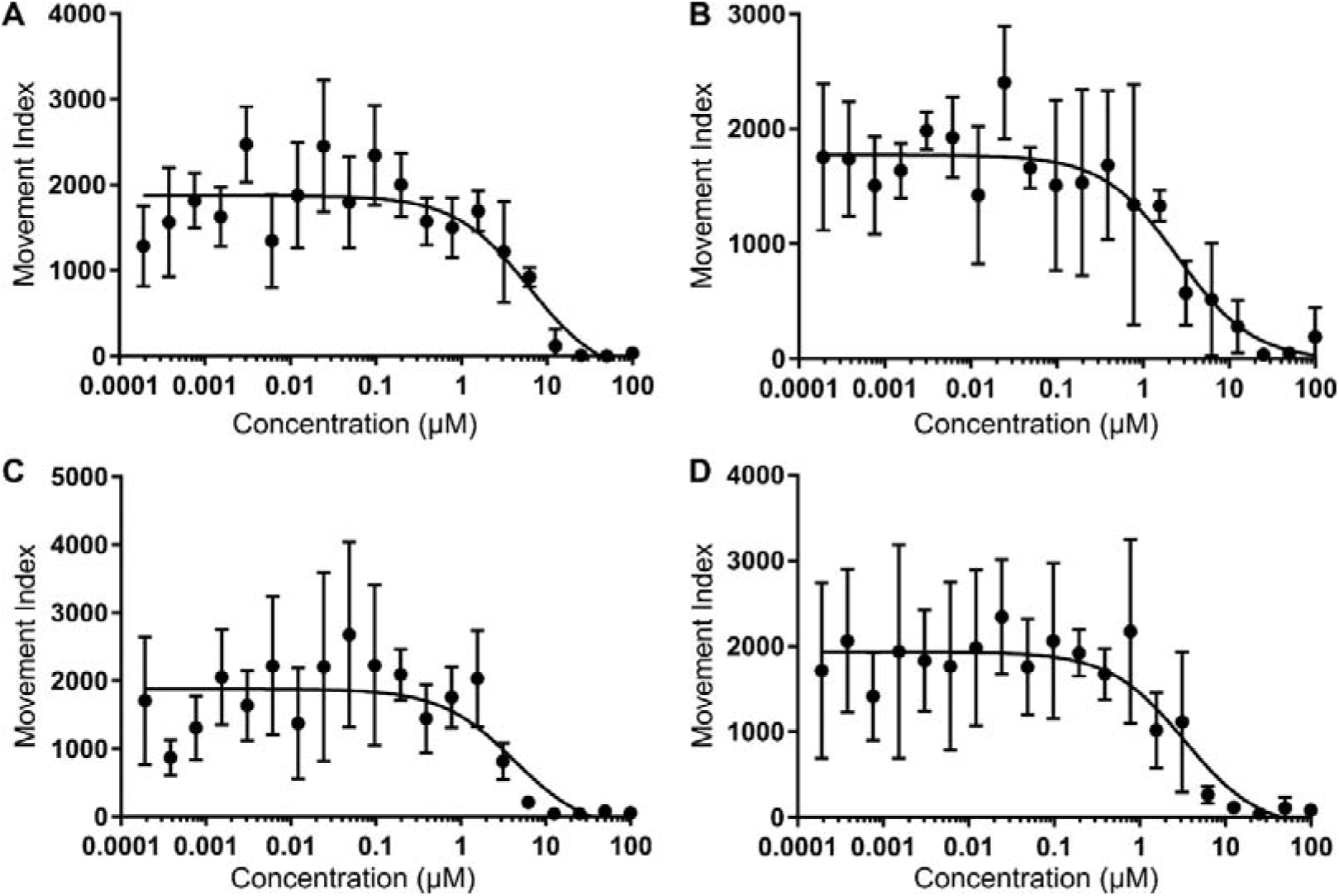
Concentration-response relationships for selected hit compounds in the *C. elegans* growth screen. (A) MMV007920 (B) MMV020152 (C) MMV652003 (D) MMV688372 MMV007920 is a benzoxazole-containing compound previously identified in a screen for agents that inhibit *Plasmodium falciparum* proliferation. The target of this compound is not known but it has been suggested that some benzoxazole compounds act on beta-tubulin (Satyendra et al., 2011). MMV020152 is an isoxazole-containing compound previously identified in a screen for compounds that inhibit *P. falciparum* growth. A number of other compounds also containing isoxazole motifs have been shown to have insecticidal activity (da Silva-Alves et al., 2013). MMV688372 is an imidazopyridine-containing compound that has been previously shown to have *in vivo* anti-trypanosomal activity (Tatipaka et al., 2014).

MMV652003 is a benzoxaborole-containing compound that has also been given the identifier AN3520 in the literature. This compound has potent activity against *Trypanosoma sp.*, both *in vitro*, and in murine models of human African trypanosomiasis (Nare et al., 2010). In this context this compound has been iteratively improved leading to the identification of the close relative SCYX-7158 (Jacobs et al., 2011), which is currently in clinical trials. The anti-trypanosomal target of this benzoxaborole class is not known (Jones et al., 2015). A simpler benzoxaborole compound, tavaborole (Fig 10B), has been approved as an anti-fungal (Elewski et al., 2015). This acts by inhibiting cytoplasmic leucyl-tRNA synthetase by forming an adduct with tRNA^Leu^ in the enzyme editing site (Rock et al., 2007). Benzoxaborole anthelmintic agents are being developed by Anacor/Eli Lilly (Akama et al., 2014; Zhang et al., 2011). Benzoxaborole compounds also show promise for other infectious diseases, including malaria, cryptosporidiosis, toxoplasmosis and tuberculosis, in each case acting via inhibition of leucyl-tRNA synthetase (Palencia et al., 2016a, 2016b; Sonoiki et al., 2016).

**Fig 10.**
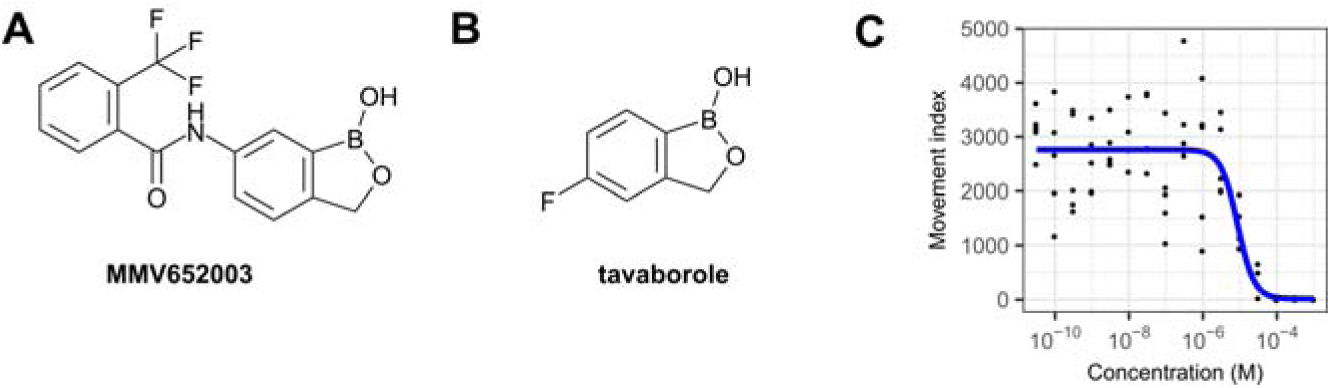
Benzoxaborole compounds as anthelmintics. Structures of (A) the screen hit MMV652003 and (B) the related benzoxaborole tavaborole, which is approved as an anti-fungal. (C) Concentration-response curve for tavaborole in the *C. elegans* growth assay.

Given the effect of MMV652003 and the similarity of this compound to the already approved drug tavaborole, we determined whether tavaborole was effective in the *C. elegans* growth assay. Tavaborole showed concentration-dependent growth inhibition (Fig 10C) with an EC_50_ of 8.6 ± 1.9 μM (Table 1). This is similar to that of the benzoxaborole screen hit, MMV652003, which has an EC_50_ of 4.3 ± 2.5 μM (Table 1). These results support the development of a benzoxaborole anti-nematode agent.

## 4. Discussion

There is an urgent need for new anthelmintics. Despite encouraging progress with MDA programs, current strategies and therapies will not achieve eradication of, for example, *T. trichiura* (Keiser J and Utzinger J, 2008; Turner et al., 2016). Furthermore, MDA, particularly with drugs that do not fully clear infection, may lead to drug resistance (Vercruysse et al., 2011). The experience from veterinary parasitology is that resistance to new anthelmintics can develop relatively rapidly after registration, causing major economic impacts and risks to global food security (Scott et al., 2013).

In this manuscript, we present the INVAPP / Paragon system which, based on imaging of whole microplates and thresholding pixel variance to determine motion, is able to quantify growth and/or motility of parasitic nematodes and the free-living nematode, *C. elegans.* A strength of this system is its high-throughput capability, typically imaging a whole plate for 5-20 seconds is sufficient to reliably quantify motion in all 96 wells. We demonstrated this effect by determining efficacy of a panel of anthelmintics in both acute motility and growth assays in *C. elegans* and of known anthelmintics in *H. contortus, T. circumcincta*, and *T. muris* assays. We further demonstrated utility of the system in a screen of small molecules for compounds that block or limit *C. elegans* growth. Current anthelmintic screens generally focus on motility reduction, as growth of parasitic nematodes can be difficult to model *in vitro*, with larvae failing to moult through their larval stages outside of the host. However anthelmintic activity *in vivo* can be much broader than inhibition of motility and thus screening compounds for their ability to inhibit *C. elegans* growth, rather than motility, represents a useful strategy to identify compounds which can subsequently be tested for growth inhibition activity *in vivo.* We used the Pathogen Box, a library of a collection of 400 diverse drug-like molecules known to be active against various neglected diseases, distributed as an open-science project by the Medicines for Malaria Venture.

Identifying the known anthelmintics mebendazole and tolfenpyrad (Preston et al., 2016b) using independent blinded screening approach serves as an important validation and supports the robustness of the screening platform. Repurposing of existing drugs for new indications is an established approach in drug discovery (Zheng et al., 2017) and is particularly valuable for neglected tropical diseases as it may reduce research and development costs and speed progress to clinical trials (Pollastri and Campbell, 2011). Auranofin has recently been shown to have activity against filarial nematode infection (Bulman et al., 2015). Our identification of auranofin as a compound that blocks *C. elegans* growth lends support to test repurposing of this compound for nematode infections. Isradipine, a safe and well-tolerated L-type calcium channel blocker, was also active in the screen. Assaying the activity of isradipine in *in vivo* models of parasitic infection is a priority and could lead to repurposing trials.

We also identified 14 compounds with previously undescribed anthelmintic activity in the *C. elegans* growth assay, belonging to a variety of chemical classes. These include benzoxazole and isoxazole compounds previously shown to have activity against *P. falciparum* (da Silva-Alves et al., 2013; Satyendra et al., 2011), and an imidazopyridine-containing compound previously shown to have *in vivo* anti-trypanosomal activity (Tatipaka et al., 2014). Another notable active compound was the benzoxaborole, MMV652003. Since the identification and successful progression into the clinic of the anti-fungal tavaborole, a number of benzoxaborole compounds have been reported to show potential for trypanosomiasis, malaria, cryptosporidiosis, toxoplasmosis and tuberculosis (Liu et al., 2014). These results support the idea that some drug chemotypes can have activity against a diversity of infectious agents. Given the costs of drug development and the limited resources available for the discovery of new drugs targeting neglected tropical diseases, it seems possible that an open source medicinal chemistry program could catalyse discovery for many different indications (Voorhis et al., 2016). Whilst preparing this manuscript we have applied the INVAPP / Paragon system to library-scale screening measuring motility of *ex vivo T. muris* adults, which led to identification of a new class of anthelmintics, the dihydrobenzoxazepinones (Partridge et al., 2017). Taken together these results demonstrate the potential for anthelmintic discovery using this system.

We focused our application of the INVAPP / Paragon system to investigating parasitic diseases. However, given its ability to determine growth and motility of *C. elegans* quickly and robustly, it could also be applied to the study of other human diseases modelled in *C. elegans*.

In conclusion, we have developed a high-throughput system for measuring the growth and/or motility of parasitic and free-living nematodes. Quantification of the activity of known anthelmintics and identification of novel chemotypes with anthelmintic activity was demonstrated, validating our approach. The system is well suited to library-scale screening of chemicals with many human and animal health applications.

## Acknowledgements

The Pathogen Box was designed and supplied by the Medicines for Malaria Venture (http://www.mmv.org). Some *C. elegans* strains were provided by the CGC, which is funded by NIH Office of Research Infrastructure Programs (P40 OD010440). F.A.P. is supported by Medical Research Council (http://www.mrc.ac.uk) grant MR/N024842/1 to D.A.L. and D.B.S. and prior to that was supported by CeBioscience Ltd and a University College London Fellowship. A.E.B. is supported by a studentship from the Rosetrees Trust. R.F. is supported by Medical Research Council (http://www.mrc.ac.uk) grant MR/N022661/1 to K.J.E. D.A.L. is supported by the NIHR UCLH Biomedical Research Centre. J.B.M. and A.A.D. are funded by the Scottish Government through the Rural and Environmental Science and Analytical Services (RESAS) Division.

## Supplementary data

**Supplementary File 1 – Results of primary and secondary screens of the Pathogen Box library using the *C. elegans* growth assay**.

